# Molecular Mechanisms of Steric Pressure Generation and Membrane Remodeling by Intrinsically Disordered Proteins

**DOI:** 10.1101/2022.05.10.491320

**Authors:** Justin R. Houser, Hyun Woo Cho, Carl C. Hayden, Noel X. Yang, Liping Wang, Eileen M. Lafer, D. Thirumalai, Jeanne C. Stachowiak

## Abstract

Cellular membranes, which are densely crowded by proteins, take on an elaborate array of highly curved shapes. Steric pressure generated by protein crowding plays a significant role in shaping membrane surfaces. It is increasingly clear that many proteins involved in membrane remodeling contain substantial regions of intrinsic disorder. These domains have large hydrodynamic radii, suggesting that they may contribute significantly to steric congestion on membrane surfaces. However, it has been unclear to what extent they are capable of generating steric pressure, owing to their conformational flexibility. To address this gap, we use a recently developed sensor based on Förster resonance energy transfer to measure steric pressure generated at membrane surfaces by the intrinsically disordered domain of the endocytic protein, AP180. We find that disordered domains generate substantial steric pressure that arises from both entropic and electrostatic components. Interestingly, this steric pressure is largely invariant with the molecular weight of the disordered domain, provided that coverage of the membrane surface is held constant. Moreover, equivalent levels of steric pressure result in equivalent degrees of membrane remodeling, regardless of protein molecular weight. This result, which is consistent with classical polymer scaling relationships for semi-dilute solutions, helps to explain the molecular and physical origins of steric pressure generation by intrinsically disordered domains. From a physiological perspective, these findings suggest that a broad range of membrane-associated disordered domains are likely to play a significant and previously unknown role in controlling membrane shape.

**Significance:** With nearly half their surfaces covered by proteins, biological membranes are highly crowded. Rapid diffusion and collision of membrane-bound proteins generates substantial steric pressure that is capable of shaping membrane surfaces. Many proteins involved in membrane remodeling, are intrinsically disordered. Having large hydrodynamic radii, disordered domains could contribute substantially to membrane crowding. However, it is unclear to what extent they are capable of generating steric pressure, owing to their conformational flexibility. Toward resolving this uncertainty, we have measured steric pressure at membrane surfaces during dynamic membrane remodeling events. Our data indicate that disordered domains generate significant steric pressure through entropic and electrostatic mechanisms, suggesting that they may constitute a critical, yet previously neglected class of membrane remodeling proteins.

## Introduction

Highly curved membrane structures are critical for cell physiology (1,2). Protein interactions on crowded cell membranes play a significant role in key membrane bending processes, such as shaping endocytic vesicles and cellular protrusions (3-5). Globular proteins with curvature-inducing motifs such as BAR domains or lattice-forming clathrin triskelia were previously thought to be the primary drivers of membrane bending (6-8). However, recent work has demonstrated that intrinsically disordered proteins are also potent drivers of membrane curvature (9,10). Specifically, the large steric bulk of disordered domains may create steric pressure on membrane surfaces, which is thought to give rise to membrane bending. While computational and analytical models have sought to predict the magnitude of this steric pressure (11-13), the ability of disordered domains to generate it has never been directly measured. As a result, the role of disordered domains in membrane remodeling remains highly debated.

Recently, we developed a FRET-based sensor for measuring steric pressure among globular proteins at membrane surfaces (14). This sensor consists of a polyethylene glycol (PEG) chain that is covalently attached at one end to the head group of a phospholipid. The free end of the PEG chain is labeled by a donor fluorophore, while lipids labeled with acceptor fluorophores reside at the membrane surface. As the membrane becomes crowded by proteins, the PEG chain is stretched such that the donor fluorophore moves farther from the membrane surface, resulting in an increase in donor lifetime. The measured donor fluorescence decay can then be used to estimate the distribution of end-to-end distances for the membrane-tethered PEG chain. Fitting these data to well-known results from the polymer physics literature provides an estimate of the change in free energy of the PEG chain due to molecular crowding, a measure of steric pressure. Importantly, our published work has already validated the ability of this sensor to accurately report the steric pressure generated by folded proteins, in agreement with crowded particle theory (14).

Here, we employ this sensor to measure the steric pressure generated by membrane-bound disordered proteins. Specifically, we examined the full-length C-terminal disordered domain of AP180 (AP180CTD-FL), one of the most abundant endocytic adaptor proteins. AP180CTD-FL has a large unstructured domain of 571 residues, which carries a high net negative charge of -32 (15). Highly charged disordered domains have the potential to generate steric pressure through both electrostatic repulsion and entropic exclusion effects (16). For example, the C-terminal domain of the neurofilament-M protein has previously been found to generate long-range repulsive forces (17), which control inter-filament spacing within axons, ultimately determining axonal diameter (18). Similarly, conformational restrictions experienced by disordered domains when they bind to membrane surfaces are thought to enable them to sense membrane curvature (16). Additionally, several recent studies have elucidated the impact of molecular crowding on the structure and conformation of disordered proteins (19-21).

Owing to the lack of secondary and tertiary structure, disordered domains are expected to be especially sensitive to the effects of molecular crowding. Indeed, work by Soranno et al. revealed polymeric behavior of disordered domains upon crowding with PEG in solution through excluded volume effects (22). Efforts have been made to study these crowding effects for synthetic polymers on lipid bilayers through both theoretical and computational approaches (23,24). However, we still lack a direct measurement of the steric forces generated by disordered domains at membrane surfaces.

In this work, we found that AP180CTD-FL generates significant steric pressure on the membrane surface, which arises from both entropic and electrostatic effects. Further, we characterized the impact of amino acid chain length on the ability of disordered proteins to generate steric pressure and drive membrane vesiculation. Our results show that steric pressure, generated by both disordered proteins and membrane-tethered polymers, depends strongly on membrane surface coverage, in agreement with classical results from polymer theory. This work provides a framework for understanding and predicting the impact of disordered domains on membrane surface pressure and shape. More broadly, our findings suggest that disordered domains, ubiquitous in membrane traffic, may constitute a broad and previously neglected class of membrane remodeling proteins.

## Results and Discussion

### The disordered domain of AP180 generates increasing steric pressure as membrane coverage increases

To determine the extent to which intrinsically disordered proteins generate steric pressure on membrane surfaces, we used AP180CTD-FL to crowd our polymer-based FRET sensor on the surfaces of small unilamellar vesicles (SUVs), which had an average diameter of about 100 nm, Figure 1A. For our FRET sensor, we used a PEG 10K chain, which was covalently attached by the manufacturer to the head group of a synthetic phospholipid, DSPE (1,2-distearoyl-sn-glycero-3-phosphoethanolamine). These PEG-conjugated lipids were incorporated into SUVs at 1 mol%. We labeled an amine group on the free end of the PEG chain with NHS-ester ATTO 488, a green fluorescent dye. ATTO 488 served as our donor fluorophore for FRET experiments. We incorporated 10 mol% of Texas Red DHPE (1,2-dihexadecanoyl-sn-glycero-3-phosphoethanolamine, triethylammonium salt) in the SUVs to serve as the FRET acceptor fluorophores. The vesicles contained 16 mol% Ni^2+^-DGS-NTA (1,2-dioleoyl-sn-glycero-3-[(N-(5-amino-1-carboxypentyl)iminodiacetic acid)succinyl]) lipids to recruit AP180CTD-FL and its variants, each of which contained an N-terminal 6x histidine tag. We varied the concentration of AP180CTD-FL in solution from 20 nM to 2 μM to increase the coverage of the membrane surface by proteins. As protein concentration increased, we measured the donor fluorescence decay, which shifted steadily toward longer times, Figure 1B. This shift suggests that crowding among AP180CTD-FL proteins drove an increase in the distribution of end-to-end distances of the polymer chains, thus limiting the number of conformations they can occupy. To model this change in conformational freedom, we fit each of the measured donor fluorescence decay curves by methods described previously (14). Briefly, we modeled the decrease in conformational entropy of the polymer chain as an effective reduction in the number of freely jointed segments within the chain. We held total chain length constant while increasing the effective length of the rigid segments that made up the chain, resulting in a decrease in the effective number of segments as shown in equation 1. Here, N’ is the effective number of segments within the crowded polymer chain, which is always less than the number of segments within the uncrowded chain, N. Similarly, r’ is the stretched segment length, which is always greater than the unstretched segment length, r. The increase in the segment length is a consequence of crowding-induced steric pressure, which mimics mechanical forces.

**Figure 1.**
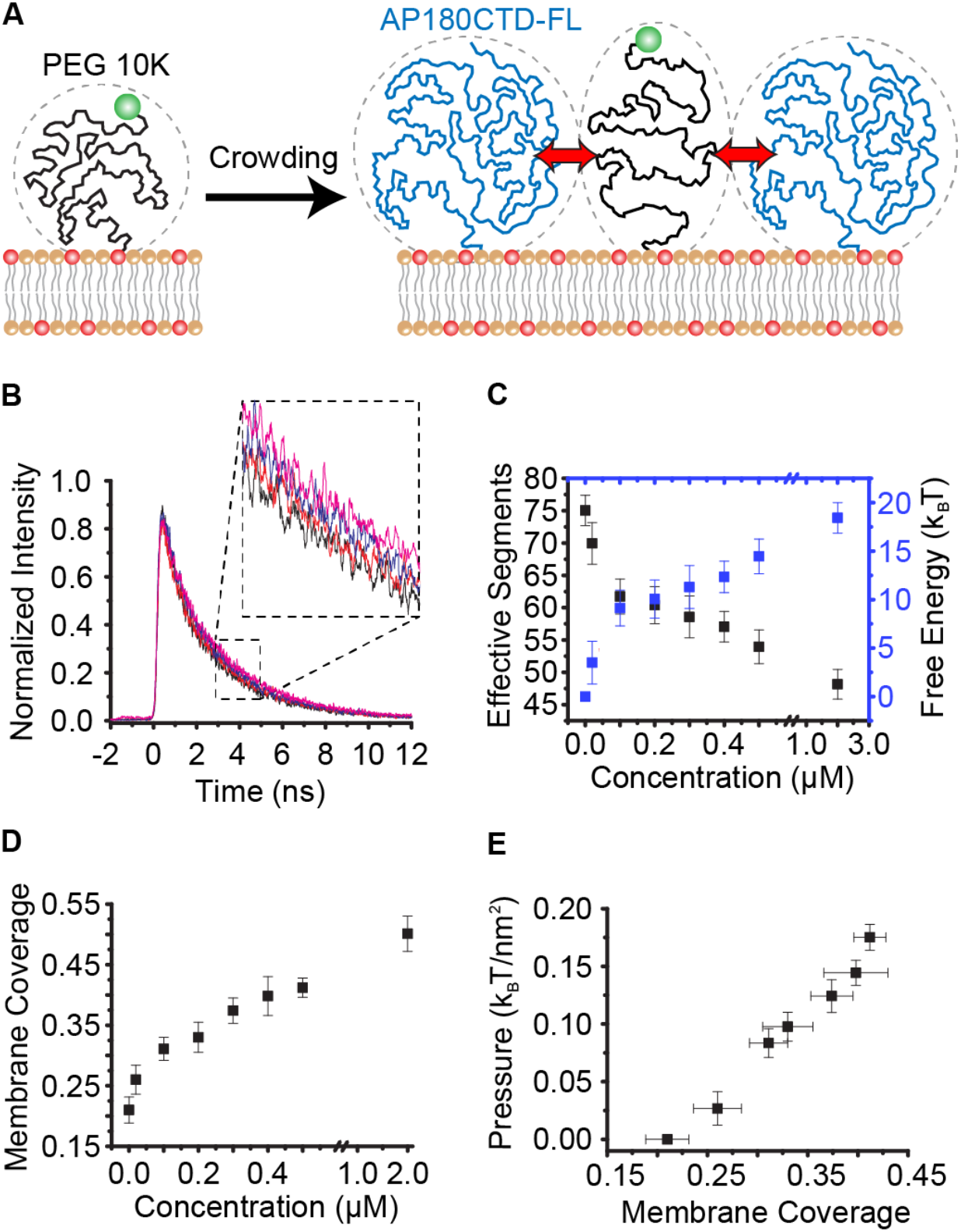
The disordered domain of AP180 generates steric pressure on membrane surfaces. The membrane composition was 72 mol% DOPC, 16 mol% Ni^2+^-DGS-NTA, 1 mol% DP-EG10-biotin, 1 mol% PEG 10K-DSPE-NH_2_, and 10 mol% Texas Red-DHPE. Vesicles were extruded to a diameter of 100 nm. The FRET donor was ATTO 488 labeled PEG 10K-DSPE-NH_2_, with the label on the free end of the polymer. The FRET acceptor was Texas Red-DHPE. (A) Schematic of the FRET sensor. PEG 10K polymers are crowded by AP180CTD-FL. (B) Donor fluorescence decay curves upon crowding with increasing concentrations of AP180CTD-FL (uncrowded black line, 200nM red line, 500nM blue line, and 2µM purple line). Curves are displayed with a moving average over a 10-point interval to better visualize the shift in decay curves. (C) Decrease in the number of polymer segments needed to fit each donor fluorescence decay curve as the concentration of AP180CTD-FL increases. The corresponding free energy is calculated for each effective segment decrease. (D) Measured fractional coverage of the membrane surface by AP180CTD-FL domains as protein concentration in solution increases. (E) Approximate steric pressure generated by AP180CTD-FL at the membrane surface with increasing fractional membrane coverage. Error bars in (C-E) are calculated as standard error of the mean.

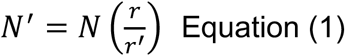

The resulting end-to-end distance of the chain increases as the effective number of segments decreases, owing to an increase in chain “stiffness”. The reduction in the effective number of segments can be used to estimate the associated reduction in chain entropy, S. Specifically, we used an analytical relationship derived by DiMarzio and McCrackin to relate the number of configurations, W, to the number of rigid segments, N, for a polymer tethered to a surface, equation 2 (25).

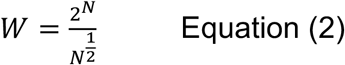

From Boltzmann’s law, the change in free energy can be estimated based on the corresponding decrease in effective segments, equation 3.

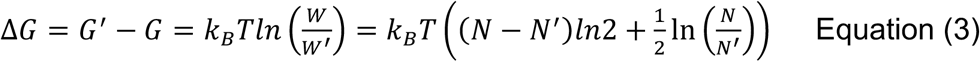

Using this analysis, we plotted the effective number of segments for each protein crowding condition along with the associated change in free energy of the polymer sensor, Figure 1C.

As the coverage of the membrane surface by AP180CTD-FL proteins increases, steric pressure is expected to increase. Therefore, we estimated the coverage of the membrane by AP180CTD-FL as a function of the protein concentration in solution. For these measurements we used a tethered vesicle assay, an approach we have previously validated, Figure 1D (10,26). In brief, biotinylated vesicles were tethered to a biotin-NeutrAvidin functionalized surface. The tethered vesicles were then incubated with a protein-containing solution and imaged, as described in the methods section. The total membrane coverage was estimated as the sum of the fraction of the membrane covered by both the polymer sensor and bound protein. The polymer sensor covered approximately 21% of the membrane surface based on our measurement of the number of PEG chains on the membrane and the projected area that each PEG chain covers, 34 nm^2^.

To estimate the steric pressure arising from protein crowding, we divided the free energy, calculated from equation 3, by the approximate area per polymer chain. In this way, the steric pressure was approximated as the change in free energy of the sensor per membrane area. We plotted the approximate steric pressure as a function of the coverage of the membrane surface by AP180CTD-FL, Figure 1E. Based on these data, we observe a clear trend of increasing steric pressure as protein coverage increases. Here we have assumed that the steric pressure restricts the conformational entropy of the disordered protein domain, which ought to result in partial stretching of its amino acid backbone. To better understand the mechanisms by which steric pressure is generated, we next sought to directly probe the straightening of the crowded disordered protein chains.

### The disordered domain of AP180 deforms measurably when crowded

Next we probed the stretching of the AP180CTD-FL protein under crowded conditions by moving the FRET sensor into the protein molecules themselves. For this purpose, we created a mutant of AP180CTD-FL that contained specific amino acid residues to be used for conjugation to the membrane surface and the donor fluorophore, respectively. At the N-terminus, we included a cysteine residue adjacent to a histidine residue, Supplemental Figure 1. Here, we used the histidine residue to chemically protect the cysteine residue during chemical conjugation of a second cysteine residue, cysteine 159, which existed natively within the AP180 C-terminal domain. Protection of the N-terminal cysteine was achieved by using cadmium ions to form a bridge between the cysteine and adjacent histidine, an established technique (27). Meanwhile, cysteine 159 was labeled with the donor fluorophore, ATTO 488-maleimide. The N-terminal cysteine was then de-protected, via removal of cadmium, and covalently conjugated to lipid vesicles through a reaction with maleimide functionalized DSPE lipids, which made up 0.5 mol% of the total membrane composition. As above, the membrane contained 10 mol% of the acceptor-labeled lipid. Once the membrane was conjugated with donor-labeled AP180CTD-FL, the surrounding space on the membrane was crowded with increasing concentrations of unlabeled AP180CTD-FL, which were recruited to the membrane surface through interactions between their N-terminal 6x-histidine tags and Ni^2+^-DGS-NTA lipids, Figure 2A. Using this system, we measured the decay of the donor fluorescence as a function of the concentration of crowders. We observed a shift towards longer donor fluorescence lifetimes, suggesting that the AP180CTD-FL protein elongated, such that the donor fluorophore moved further from the acceptor labeled membrane surface as the membrane became crowded, Figure 2B. To approximate the distance that AP180CTD-FL was stretched by crowding, we fit a single component exponential decay to the measured donor decay curve of the sensor, Supplemental Figure 2. We used the lifetime, *τ*_*DA*_, of this exponential to calculate the FRET efficiency, E, relative to the unquenched lifetime of ATTO 488, *τ*_*D*_, in the absence of acceptors according to equation 4.

**Figure 2.**
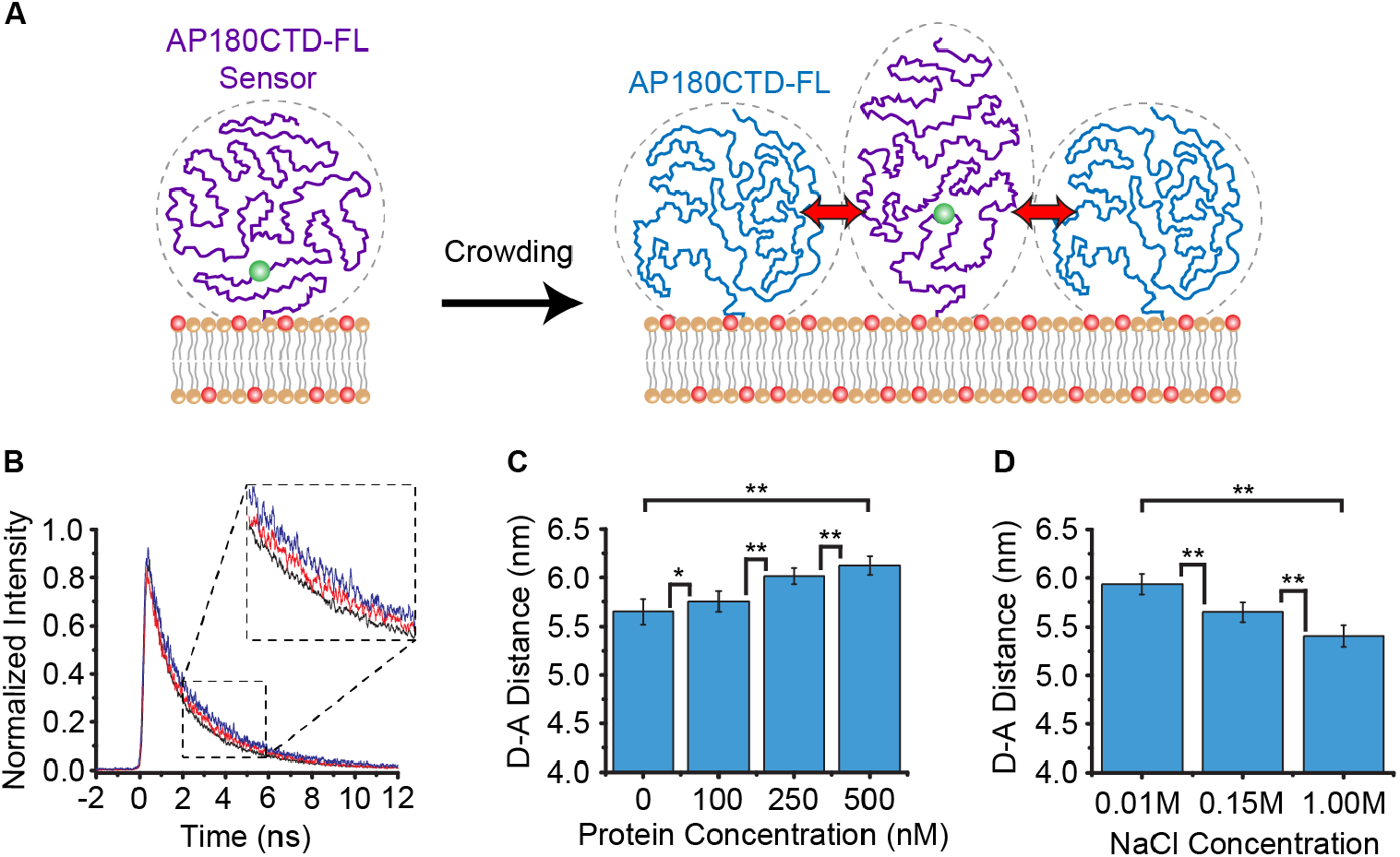
AP180CTD-FL deforms in response to changes in crowding and ionic strength. The membrane composition was 72.5 mol% DOPC, 16 mol% Ni^2+^-DGS-NTA, 1 mol% DP-EG10-biotin, 0.5 mol% DSPE-MAL, and 10 mol% Texas Red-DHPE, extruded to a diameter of 100 nm. (A) Schematic of the FRET sensor embedded within AP180CTD-FL. Crowding by other unlabeled AP180CTD-FL domains occurs with increasing membrane coverage. (B) Donor fluorescence decay curves upon crowding with increasing concentration of AP180CTD-FL (uncrowded black line, 250 nM red line, 500 nM blue line). (C) Increase in the characteristic donor-acceptor distance of AP180CTD-FL as protein concentration in solution increased. (D) Decrease in the average donor-acceptor distance of AP180CTD-FL as the ionic strength of the solution increased. Error bars in (C) and (D) are calculated as standard error of the mean. P values in (C) and (D) were calculated using unpaired, two-tailed Student’s t tests. *P < 0.05, **P < 0.01.

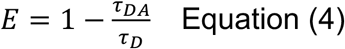

We used the calculated FRET efficiencies from equation 4 to calculate the characteristic donor-acceptor distance, R_DA_, in each AP180CTD-FL crowding condition, Figure 2C. The donor-acceptor distance is related to the FRET efficiency through the Förster radius, R_0_, of the donor-acceptor FRET pair, approximately 5.4 nm for ATTO 488 and Texas Red, equation 5.

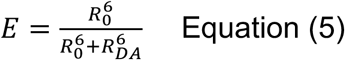

From these crowding experiments, we observed an increase of approximately 0.48 nm from an uncrowded environment to the most crowded condition. This relatively small deformation demonstrates that AP180CTD-FL does not undergo the dramatic conformational changes required to generate a polymer-like brush on the membrane surface (28,29). Instead, AP180CTD-FL likely generates steric pressure using a combination of entropic and electrostatic mechanisms in the semi-dilute regime of polymer crowding (30). Specifically, the substantial net negative charge of AP180CTD-FL could contribute to electrostatic repulsion between chains, leading to an enhanced crowding effect at high crowding densities. In accord with this hypothesis, we observed a decrease of 0.52 nm in the donor-acceptor distance of AP180CTD-FL chains as we increased the ionic strength of the solution from 10 mM NaCl to 1M NaCl, Figure 2D. Observing this sensitivity to ionic strength of the solution, we next asked to what extent do electrostatic interactions play a role in generating steric pressure on the membrane surface?

### Electrostatic interactions contribute to steric pressure generation by the disordered domain of AP180

The C-terminal domain of AP180 has a relatively high net negative charge of -32 (15). In a previous work, we observed a substantial decrease in AP180’s hydrodynamic radius in solutions of high ionic strength, where electrostatic screening is strong (16). This result is consistent with our finding in Figure 2 that donor fluorophores placed within AP180 come closer to the membrane surface when AP180’s net charge is screened under conditions of high ionic strength. Therefore, we hypothesized that AP180CTD-FL’s ability to generate steric pressure on the membrane could be enhanced by electrostatic repulsion, Figure 3A. Using the same vesicle composition as described in Figure 1, we repeated our FRET experiment at ionic strengths of 10 mM, 150 mM, and 1M NaCl, while crowding the membrane with a fixed concentration of 500 nM AP180CTD-FL, Figure 3B. As ionic strength increased, the donor fluorescence decay shifted toward shorter times, such that the effective number of segments used to fit the decay curves increased with increasing salt concentration, when protein concentration in solution was held constant, Figure 3C. We next measured the impact of electrostatic screening on coverage and found that it increased with increasing ionic strength, likely due to a decrease in repulsion between AP180CTD-FL and the membrane surface, which contains negatively charge Ni^2+^-DGS-NTA lipids, Figure 3D. Notably, calculating the coverage requires that measured values of protein density per membrane area be multiplied by the projected area of AP180CTD-FL on the membrane surface, which depends on the protein’s hydrodynamic radius. Since changing the ionic strength of the buffer also changes the hydrodynamic radius of the protein, we used values for the hydrodynamic radius at each salt concentration previously determined by Zeno et al. (16). Combining data in Figures 3C and 3D over a range of protein concentrations in solution, we plotted steric pressure as a function of membrane coverage for AP180CTD-FL at ionic strengths of 10 mM, 150mM, and 1M NaCl. These data demonstrate that AP180CTD-FL generates higher pressures at lower salt concentrations, where electrostatic screening is weakest, such that electrostatic repulsion makes its strongest contribution. In particular, the steric pressure at 10 mM NaCl is more than twice that at 1M NaCl. These results demonstrate that electrostatic repulsion between negatively charged disordered domains makes a substantial contribution to steric pressure generation at membrane surfaces. We next asked, to what extent does the length of the disordered domain impact steric pressure?

**Figure 3.**
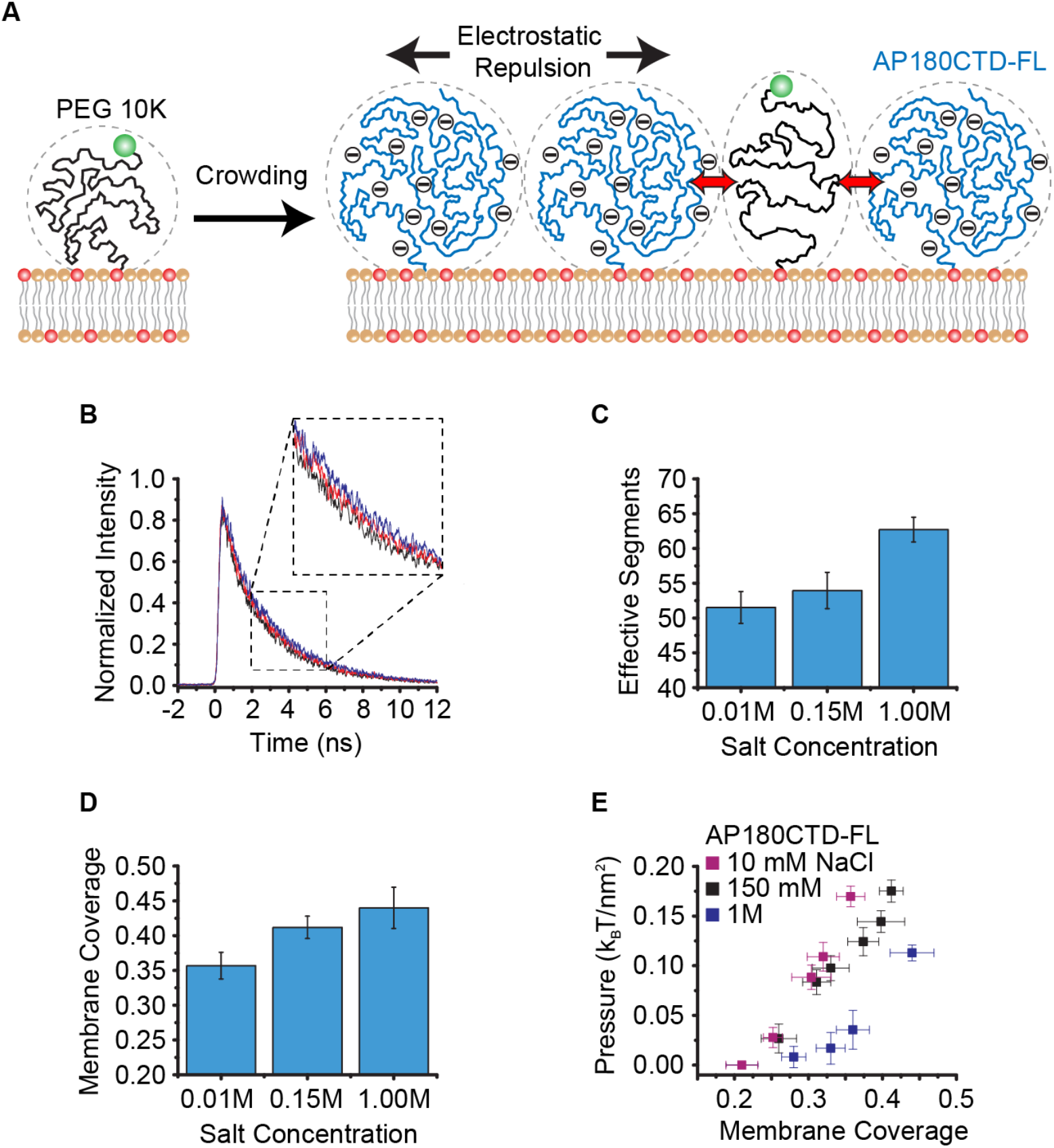
Ionic strength impacts steric pressure generated by AP180CTD-FL. The membrane composition was 72 mol% DOPC, 16 mol% Ni^2+^-DGS-NTA, 1 mol% DP-EG10-biotin, 1 mol% PEG 10K-DSPE-NH_2_, and 10 mol% Texas Red-DHPE. Vesicles were extruded to a diameter of 100 nm. (A) Schematic of the electrostatic interactions between AP180CTD-FL domains and the impact on crowding the PEG sensor. (B) Donor fluorescence decay curves with decreasing NaCl concentration (1 M NaCl black line, 150 mM NaCl red line, and 10 mM NaCl blue line). All curves shown are at a constant protein concentration of 500 nM AP180CTD-FL. Curves are displayed with a moving average over a 10-point interval to better visualize the shift in decay curves. (C) Increase in number of polymer segments needed to fit each donor fluorescence decay curve at a constant protein concentration (500 nM) as the NaCl concentration in solution increased. (D) Measured fractional coverage of the membrane surface by AP180CTD-FL at a constant protein concentration (500 nM) as the NaCl concentration in solution increased. (E) Approximate steric pressure generated by AP180CTD-FL at the membrane surface with increasing NaCl concentration. Error bars in (C) are calculated as standard deviation of the mean. Error bars in (D) are calculated as standard error of the mean.

### The length of the disordered domain does not significantly impact steric pressure generation at constant coverage

To evaluate the effect of protein chain length on steric pressure generated at the membrane surface, we examined an N-terminal truncation of the C-terminal domain of AP180 (AP180CTD-1/3), which consisted of approximately the first third (190 amino acids) of the full-length protein, Supplemental Figure 3. Using the same methods and vesicle composition as described above, we used the PEG 10K sensor to examine steric pressure generation by AP180CTD-1/3 on crowded membrane surfaces, Figure 4A. As the concentration of AP180CTD-1/3 increased in solution, we observed that the donor fluorescence decay curve shifted towards longer times, similar to our results with AP180CTD-FL, Figure 4B. Using the same approach described above, we evaluated the impact of AP180CTD-1/3 on the effective number of segments in the PEG 10K sensor, and used the change in effective segment number to estimate the change in free energy per membrane area upon crowding by AP180CTD-1/3, Figure 4C. Additionally, we measured the coverage of AP180CTD-1/3 on the membrane surface using the tethered vesicle assay, Figure 4D. Using these data, we constructed a pressure-coverage curve for AP180CTD-1/3, which is plotted alongside the curve for AP180CTD-FL, Figure 4E. Interestingly, we observed that AP180CTD-1/3 generated only slightly less pressure than AP180CTD-FL. Further, when we reduced the contribution of electrostatic interactions by increasing the ionic strength of the solution to 1M NaCl, the pressure versus coverage curves for AP180CTD-1/3 and AP180CTD-FL collapsed onto approximately the same curve, Figure 4F.

**Figure 4.**
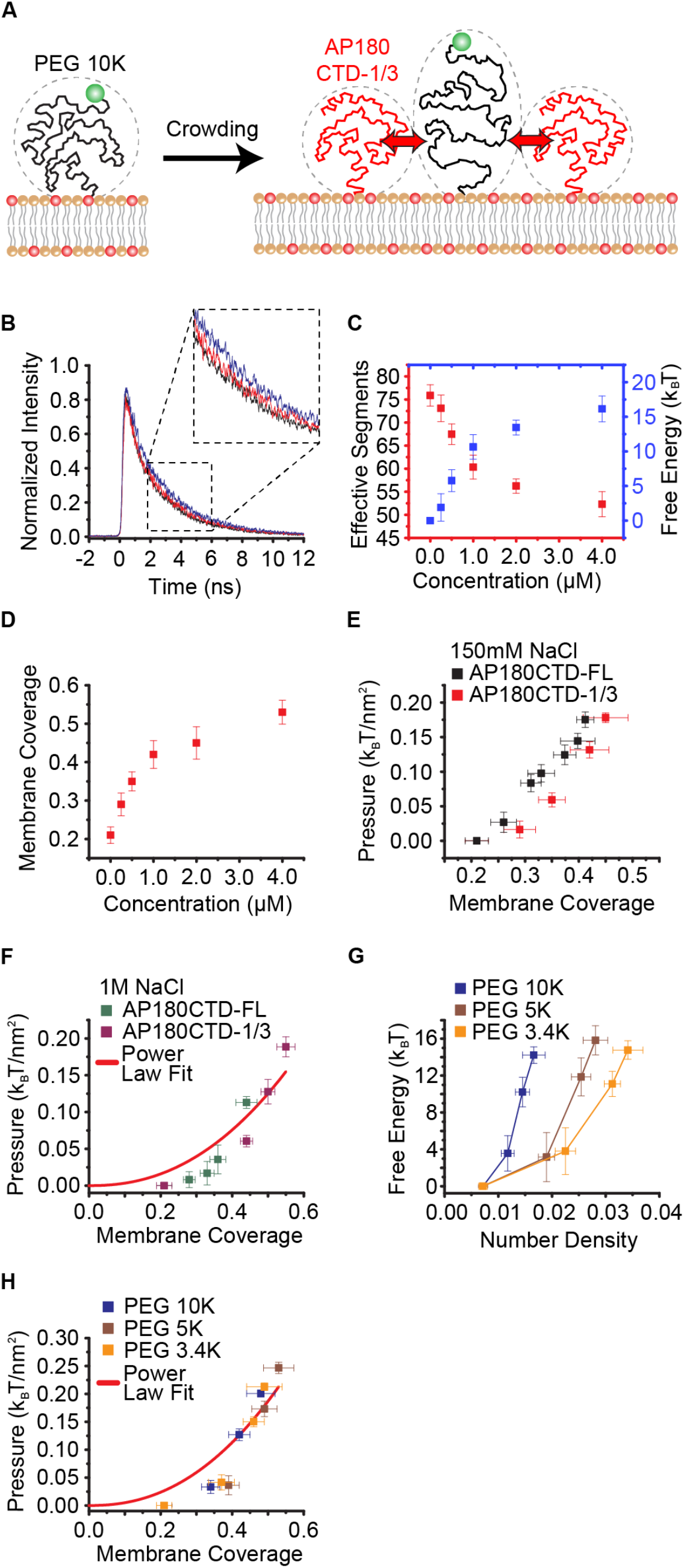
The chain length of AP180CTD and PEG does not significantly impact steric pressure generated at the membrane surface at constant coverage. For AP180CTD crowding experiments, the membrane composition was 72 mol% DOPC, 16 mol% Ni^2+^-DGS-NTA, 1 mol% DP-EG10-biotin, 1 mol% PEG 10K-DSPE-NH_2_, and 10 mol% Texas Red-DHPE. Vesicles were extruded to a diameter of 100 nm. For PEG crowding experiments, the membrane composition was 83-85.75 mol% DOPC, 1 mol% DP-EG10-biotin, 1 mol% PEG 10K-DSPE-NH_2_, 2.25-5 mol% PEG 3.4K-DSPE-SH_2_, PEG 5K-DSPE-SH_2_, or PEG 10K-DSPE-SH_2_ and 10 mol% Texas Red-DHPE. (A) Schematic of AP180CTD-1/3 crowding the PEG sensor. (B) Donor fluorescence decay curves upon crowding with increasing concentration of AP180CTD-1/3 (uncrowded black line, 500nM red line, 2 μM blue line). Curves are displayed with a moving average over a 10-point interval to better visualize the shift in decay curves. (C) Decrease in the number of polymer segments needed to fit each donor fluorescence decay curve as the concentration of AP180CTD-1/3 increased. The corresponding free energy was calculated for each effective segment decrease. (D) Measured fractional coverage of the membrane surface by AP180CTD-1/3 domains as protein concentration in solution increased. (E) Approximate steric pressure generated by AP180CTD-1/3 compared to AP180CTD-FL as membrane coverage increased with 150 mM NaCl in solution. (F) Approximate steric pressure generated by AP180CTD-1/3 compared to AP180CTD-FL as membrane coverage increased with 1M NaCl in solution. All data points were grouped together and fit (red line) by a power law, y = Ax^9/4^, where A was a free fitting parameter. (G) Calculated free energy of PEG crowding for different sizes of PEG as the number density of PEG chains on the membrane surface increased. (H) Approximate steric pressure associated with crowding different sizes of PEG as the fractional membrane coverage of PEG chains increased. All data points were grouped together and fit (red line) by a power law, y = Ax^9/4^, where A was a free fitting parameter. Error bars in (C-H) were calculated as standard error of the mean.

Notably, when rigid particles are crowded, steric pressure increases with the inverse particle size, when coverage is held constant. Therefore, the lack of a clear dependence on protein molecular weight in our results likely arises from the flexible, polymer-like nature of intrinsically disordered proteins. Why would steric pressure generation be essentially independent of chain length at constant membrane coverage for disordered proteins? First, it is important to note that the coverage of the membrane surface by proteins in our measurements is in the semi-dilute regime, approximately 20-50% (30). In this regime, adjacent polymers begin to contact one another, yet the polymer chains are not substantially reconfigured or stretched by these interactions, as demonstrated by the data in Figure 2. Therefore, for long chains, monomers have a similar probability of interacting, whether they are part of the same chain or are derived from two adjacent chains. In this way, interactions among the chains depends more on the overall monomer concentration than on the lengths of the individual chains. Under these assumptions, the Des Cloiseaux law, a well-known result from polymer physics, describes the thermodynamics of polymers (30,32). Specifically, it predicts that all thermodynamic properties of polymers in the semi-dilute regime must reach a limit that depends on monomer concentration, but is independent of the degree of polymerization, i.e. the length of the chains. It follows that in this semi-dilute regime, the local free energy is controlled mainly by the monomer concentration in the solution. Thus, the osmotic pressure, which is analogous to the steric pressure we measure, can be described by equation 6, where Π is osmotic pressure, a is segment length, T is temperature, and Φ is coverage.

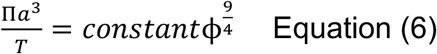

Equation 6 predicts that, for idealized chains at a fixed coverage, pressure should be independent of chain length, increasing approximately as the 9/4 power of coverage. This trend is in reasonable agreement with the pressure versus coverage curves for AP180CTD-FL and AP180CTD-1/3 at 1M NaCl, Figure 4F.

In order to further investigate the impact of chain length on steric pressure generation, we also examined crowding by membrane-tethered polyethylene glycol (PEG) chains. As synthetic homopolymers, PEG chains are perhaps better approximated as idealized chains, as assumed by the Des Cloiseaux law. For these experiments, we used a similar vesicle composition as described above with 1 mol% donor-labeled PEG 10K as our sensor and 10 mol% Texas Red-DHPE to quench the donor. However, instead of incorporating Ni^2+^-DGS-NTA lipids into the vesicles, we used an increasing mol% from 2.25% to 5% of PEGylated DSPE lipids to crowd the membrane surface. We performed our FRET experiments as described above and calculated the corresponding steric pressure for each PEG crowding condition. In order to approximate the number density of crowding PEGs on the membrane surface, we again used the tethered vesicle assay, see methods. Figure 4G plots steric pressure as a function of the number density of PEG chains on the membrane surface. When number density was held constant, steric pressure increased with increasing length of the PEG chain, owing to the higher coverage of the larger chains. However, when steric pressure was plotted as a function of membrane coverage, all the curves collapsed onto a single curve, Figure 4H, as predicted by the Des Cloiseaux scaling. Here the hydrodynamic radii of the PEG chains are used to estimate coverage, based on the measured values of number density, see methods. We surmise that Equation 6 provides a reasonable qualitative fit to these data, given that the resolution of our sensor is limited at low pressures (14). Taken together, these data demonstrate that steric pressure generated by crowded polymer-like molecules on membrane surfaces has a weak dependence on chain length, in the semi-dilute regime for long chains, in approximate agreement with the Des Cloiseaux scaling. Notably, the scaling of steric pressure with chain length is likely to become more complicated when a broader range of lengths is considered, and the Des Cloiseaux scaling may not accurately describe the full range of physiologically relevant disordered protein chains.

### The disordered domain of AP180 drives membrane vesiculation by a mechanism that is largely independent of protein chain length

We have previously shown that intrinsically disordered protein domains are capable of generating sufficient steric pressure to drive vesiculation, or breakup of membrane vesicles to form smaller “daughter” vesicles (33). Here we asked how the ability to drive vesiculation might depend on the length of the disordered protein domain. Specifically, we compared membrane vesiculation by AP180CTD-FL and AP180CTD-1/3, Figure 5A. We used a quantitative fluorescence approach to measure the distribution of vesicle diameters following protein exposure, a technique that we have reported previously, Figure 5B (26,33). Briefly, vesicles were incubated with either AP180CTD-FL or AP180CTD-1/3 at concentrations varying from 20 nM to 4 µM for 1 hour at room temperature. After incubation, the resulting vesiculation products were tethered to a biotin-NeutrAvidin functionalized coverslip. The distribution of vesicle diameters was then determined by comparing the fluorescence intensity distributions before and after exposure of the vesicles to proteins. Here an approximate conversion between vesicle brightness and vesicle diameter was determined by comparing the brightness distribution of vesicles prior to protein exposure, with the distribution of diameters, measured using dynamic light scattering. Additional details can be found in the methods section and in our previously published work (26,33). We observed a steady decrease in vesicle diameter with increasing concentration of protein in solution, Figures 5C, 5D. Using the previously measured coverage of each protein on the membrane surface as a function of its concentration in solution, Figures 1D and 4D, we plotted vesicle diameter as a function of membrane coverage by proteins, Figure 5E, and as a function of steric pressure, Figure 5F. As the fractional membrane coverage of AP180CTD-FL and AP180CTD-1/3 increased to about 50%, we observed a decrease in vesicle diameter to around 35 nm. Importantly, when we compare AP180CTD-FL and AP180CTD-1/3 at equal membrane coverage, similar amounts of steric pressure are generated and similar trends of membrane fission to smaller diameters occur. This result demonstrates that AP180CTD drives membrane fission by a mechanism that is largely independent of its chain length, when membrane coverage is held constant.

**Figure 5.**
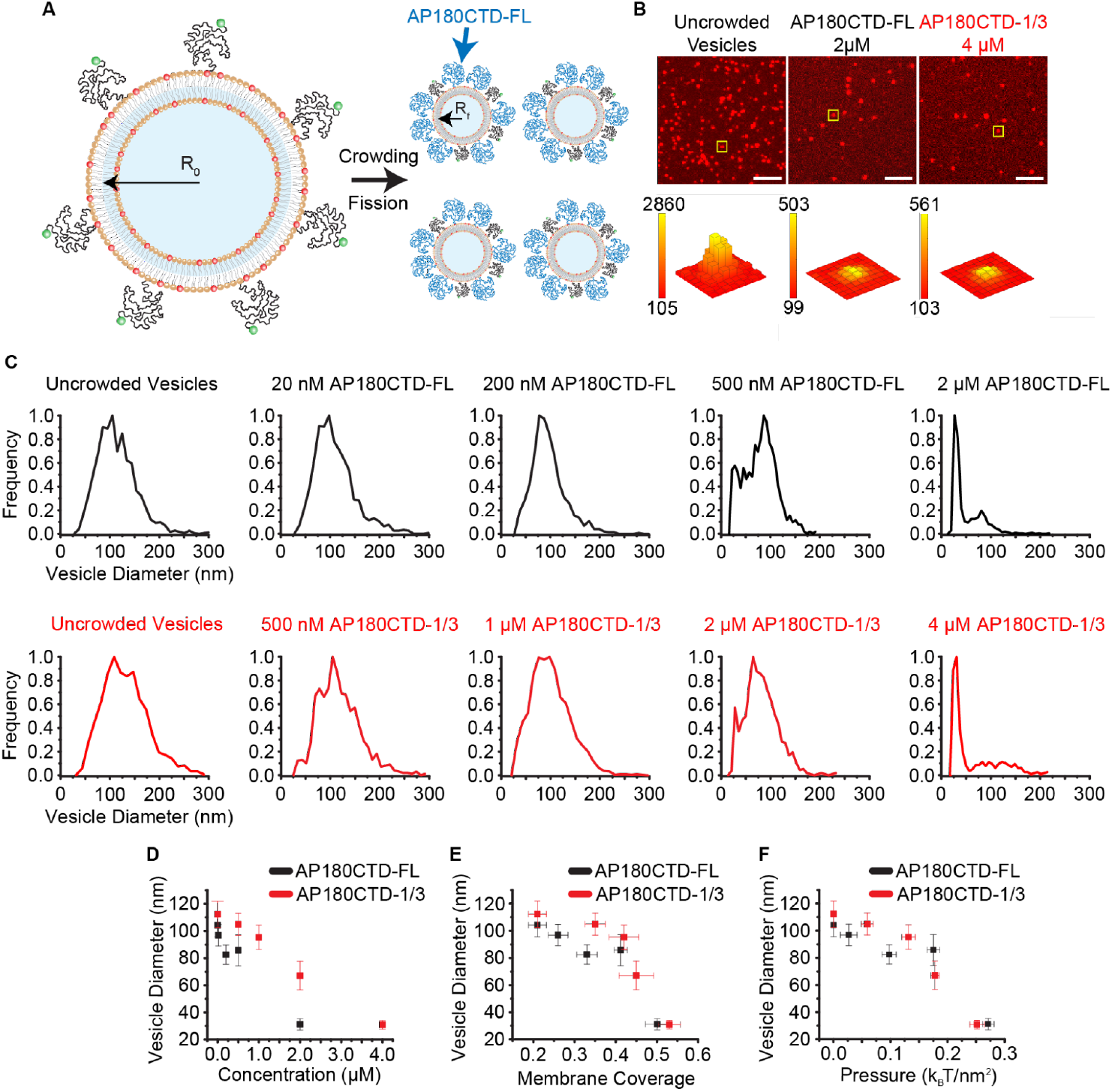
Disordered domains that achieve similar pressure on the membrane surface drive equivalent levels of membrane fission. The membrane composition was 81.9 mol% DOPC, 16 mol% Ni^2+^-DGS-NTA, 1 mol% DP-EG10-biotin, 1 mol% PEG 10K-DSPE-NH_2_, and 0.1 mol% Texas Red-DHPE. Vesicles were extruded to an initial diameter of 100 nm. (A)Schematic of membrane crowding by disordered proteins and resulting fission. Vesicles started at a radius of R_0_ and underwent fission into several smaller vesicles, ending with a radius of R_f_. (B) Fluorescent images of lipid vesicles acquired by spinning disk confocal microscopy. Vesicles were crowded by AP180CTD-FL and AP180CTD-1/3 and underwent fission, resulting in a significantly decreased fluorescence intensity. An example puncta is highlighted by a yellow box to show the fluorescence distribution. Scale bars are 5 µm. (C) Distributions of vesicle diameters as the protein concentration of AP180CTD-FL (black) or AP180CTD-1/3 (red) was increased. Distributions are binned histograms of data analyzed from the tethered vesicle assay. (D) Vesicle diameter as the concentration of AP180CTD-FL or AP180CTD-1/3 in solution increased. (E)Vesicle diameter as membrane coverage by AP180CTD-FL or AP180CTD-1/3 increased. (F) Vesicle diameter as steric pressure generated by AP180CTD-FL or AP180CTD-1/3 increased. Error bars in (D-F) were calculated as standard error of the mean.

## Conclusions

In this work, we demonstrate the first measurement of steric pressure generated by disordered proteins bound to membrane surfaces. Using this measurement, we have shown that the high net negative charge of AP180CTD-FL contributes to its ability to generate steric pressure on membrane surfaces. Further, AP180CTD-FL elongates slightly when crowded by sufficient membrane coverage. This slight elongation suggests that crowding occurs in the semi-dilute regime of coverage on the membrane surface. From these results, we hypothesized that both the electrostatic repulsion between chains and steric effects of crowding led to pressure generation on the membrane surface. Interestingly, we found that pressure generated by disordered proteins was largely independent of the amino acid chain length. We confirmed these results by examining crowded PEG chains on membrane surfaces, which provided a model of an ideal homopolymer. Interestingly, the Des Cloiseaux scaling for free polymers in the semi-dilute regime offers a reasonable explanation for the relative independence of steric pressure on chain length at constant membrane coverage. Finally, we investigated the impact of steric pressure on the physical processes of membrane vesiculation. Here we found that the capacity for membrane vesiculation was largely independent of disordered protein length, when membrane coverage was held constant. These results demonstrate a general polymer-like crowding mechanism for generating steric pressure and ultimately shaping and vesiculating membrane surfaces.

Our work contributes to a physical understanding of how disordered domains generate steric pressure at membrane surfaces. Future work has the potential to develop a predictive understanding of the role of intrinsically disordered proteins and steric pressure in membrane bending and vesiculation processes in the cell. In particular, more than 30% of proteins involved in clathrin-mediated endocytosis contain long disordered regions of at least 100 amino acids (34). Our work suggests that any of these regions could contribute significantly to steric pressure generation and membrane bending, suggesting that the “proteome” of membrane remodeling may be much larger than previously imagined.

## Materials and Methods

### Materials

DOPC (1,2-dioleoyl-sn-glycero-3-phosphocholine) and Ni^2+^-DGS-NTA (1,2-dioleoyl-sn-glycero-3-[(N-(5-amino-1-carboxypentyl)iminodiacetic acid)succinyl] nickel salt) were purchased from Avanti Polar Lipids, Inc. DP-EG10-biotin (dipalmitoyl-decaethylene-glycol-biotin) was generously provided by Darryl Sasaki from Sandia National Laboratories, Livermore, CA. ATTO 488 NHS-ester was purchased from ATTO-TEC GmbH. NeutrAvidin and Zeba spin desalting columns were purchased from Thermo Fisher Scientific. TCEP (Tris(2-carboxyethyl) phosphine hydrochloride), EDTA-free protease inhibitor tablets, PLL (poly-L-lysine), Texas Red DHPE (1,2-dihexadecanoyl-sn-glycero-3-phosphoethanolamine, triethylammonium salt), DMSO (dimethyl sulfoxide), cadmium chloride, EDTA (Ethylenediaminetetraacetic acid), and Thrombin CleaveCleave Kit were purchased from Sigma-Aldrich. Sodium chloride, potassium chloride, sodium bicarbonate, HEPES (4-2(2-hydroxymethyl)-1-piparazineethanesulphonic acid), IPTG (isopropyl-β-D-thiogalactopyranoside), and Triton X-100 were purchased from Fisher Scientific. Amine reactive PEG (mPEG-Succinimidyl Valerate MW 5000) and PEG-biotin (Biotin-PEG SVA MW 5000) were purchased from Laysan Bio, Inc. Amicon Ultra centrifugal filter units were purchased from MilliporeSigma. PEG 3.4K, 5K, and 10K DSPE (1,2-distearoyl-sn-glycero-3-phosphoethanolamine) SH_2_, PEG 10K DSPE-NH_2_, and DSPE-maleimide were purchased from Nanosoft Polymers. Centri-spin-20 size exclusion columns were purchased from Princeton Separations. All reagents were used without additional purification.

## Methods

### Quantitative FRET Measurements by TCSPC

Coverslips with 1.5 thickness (13-16 μm) were cleaned by a heated acid, base, and hydrogen peroxide washing technique. To passivate the surface of the coverslip before imaging, we created a supported lipid bilayer in small silicone wells on the coverslip. This bilayer prevented significant lipid or protein binding to the coverslip. We then added the lipid vesicle mixture to the wells and proceeded to collect fluorescence lifetime data. A homebuilt point-scanning confocal microscope was used to acquire all lifetime data. A 486 nm picosecond pulsed diode laser at a repetition rate of 50 MHz was used to excite the donor fluorophore (ATTO 488). The laser beam was focused 5 μm above the surface of the glass coverslip using a piezoelectric stage (Mad City Labs). Sample fluorescence was collected by a 100x magnification, 1.4 NA microscope objective, focused through a pinhole onto a GaAsP photomultiplier tube (Hamamatsu). Sample fluorescence was passed through a 511 nm bandpass emission filter with a 20 nm bandwidth before arriving at the photomultiplier tube. The photomultiplier tube output pulses were amplified and sent to specialized electronics for time-correlated single-photon counting (Becker and Hickl). Donor fluorescence decay curves were collected for 100-300 seconds, and 9-30 decay curves were collected across at least two independent samples per data point.

### Lipid vesicle preparation for FRET

Small unilamellar vesicles (SUVs) were prepared, which incorporated 1 mol% of PEG 10K, covalently attached to 1,2-distearoyl-sn-glycero-3-phosphoethanolamine (DSPE). The free ends of the PEG chains displayed an amine group, which we used to attach the donor fluorophore to the end of the PEG chain (Nanosoft Polymers), as described below. Texas Red covalently attached to 1,2-Dihexadecanoyl-sn-Glycero-3-Phosphoethanolamine (DHPE) served as the acceptor fluorophore in the FRET system, and was included in the lipid composition at 10 mol%. 16 mol% of 1,2-dioleoyl-sn glycero-3-[(N-(5-amino-1-carboxypentyl)iminodiacetic acid)succinyl] nickel salt (Ni^2+^-DGS-NTA) was used to bind 6x histidine tagged AP180CTD-FL and AP180CTD-1/3 proteins in all experiments. 1 mol% dipalmitoyl-decaethylene-glycolbiotin (DP-EG10-biotin) was incorporated into the vesicles for all tethered vesicle experiments. The remainder of each vesicle composition consisted of 1,2-dioleoyl-sn-glycero-3-phosphocholine (DOPC). The lipids were mixed in a clean glass test tube, the solvent was evaporated, and the lipid film was further dried under vacuum overnight. The dried lipid film was hydrated in 20 mM sodium bicarbonate, 150 mM KCl, pH 8.2 at room temperature for 15 minutes. The lipid mixture was then extruded through a filter with 100 nm pores. The extruded vesicles were then incubated with the donor dye (100 μM of NHS-ester functionalized ATTO 488) for 1 hour at room temperature. We then successively washed the excess dye out from the mixture using 100K Amicon spin filter units (MilliporeSigma cat#UFC510024). In order to minimize background levels of free dye in solution, the solution surrounding the vesicle was effectively diluted by 10^14^ fold during the washing step, where the vesicles were finally left in the experimental buffer consisting of 25 mM Hepes, 150 mM NaCl, pH 7.35.

### Protein Expression and Purification

AP180CTD and related constructs were each expressed as fusion proteins with an N-terminal GST tag for increased stability. GST was subsequently removed by thrombin cleavage. *E. coli* BL21 competent cells (NEB Cat#C2530), transformed with a plasmid encoding either AP180CTD-FL or AP180CTD-1/3, were grown at 30°C until the suspension reached an OD600 of 0.8. Protein expression was induced with 1 mM IPTG for 6-8 h at 30 °C. Cells were pelleted and stored at −80 °C. The whole protein purification was performed at 4 °C. Cells were resuspended in lysis buffer [0.5 M Tris pH 8.0, 5 mM EDTA, 5 v/v % glycerol, 5 mM DTT, Roche protease inhibitor cocktail (Roche cat#05056489001) pellets 1/50 mL] + 1 v/v% Triton X-100. The suspension was sonicated on ice (4 × 2000 J). The mixture was clarified by centrifugation at 26581g for 25 min, and the supernatant was chromatographed on a glutathione Sepharose column. The column was washed with 10 column volumes of lysis buffer + 0.2% Triton X-100 followed by another 5 column volumes of lysis buffer with no added Triton X-100. GST-containing proteins were eluted from the column with 15 mM glutathione in lysis buffer and exchanged into 50 mM Tris (pH 8.0), 10 mM CaCl_2_, 150mM NaCl, 5mM EDTA using a Zeba desalting column (Thermo Scientific cat#89891). GST was cleaved using the Thrombin CleanCleave kit (Sigma-Aldrich cat#RECOMT) for 14 h at 4 °C with gentle rocking. The cleaved GST and any uncut fusion protein was removed with another glutathione Sepharose column. Purified protein was then concentrated using an Amicon Ultra-15 centrifugal filter (MilliporeSigma cat#UFC903024) and stored as liquid nitrogen pellets at −80 °C.

### Protein Labeling

ATTO-488 NHS-ester was dissolved in dimethyl sulfoxide (DMSO) at a concentration of 10 mM and stored at -80 °C. Primary amines within the proteins were then labeled in buffer consisting of in 20 mM sodium bicarbonate, 150 mM KCl, pH 8.2. Protein concentration varied from 20 to 150 μM. The dye solution in DMSO was added to the protein solution at a stoichiometric ratio of 3 dye molecules per protein molecule and allowed to react for 30 minutes at room temperature. The total amount of DMSO present never exceeded 1 v/v %. Labeling ratios for the proteins varied from 0.5 to 2.1 dyes per protein. Unreacted dye was removed using Centri-Spin-20 size exclusion columns (Princeton Separations). Protein and dye concentrations were measured using UV−vis spectroscopy, and labeled proteins were stored at −80 °C.

### Protected Cysteine Labeling of AP180CTD-FL

In order to label AP180CTD-FL at a specific site, we introduced N-terminal cysteine (C8) and histidine (H7) residues directly adjacent to each other in the linker region of AP180CTD-FL, Supplemental Figure 1. We used a previously published technique of labeling site specific cysteine residues by reversible cadmium metal protection (27). The cysteine metal affinity for cadmium is enhanced by an adjacent histidine residue, therefore making the cysteine inaccessible to standard maleimide labeling techniques. In contrast, the second native cysteine (C159) has no histidine adjacent to it, and therefore does not bind cadmium, making it accessible for maleimide labeling. We first protected the N-terminal cysteine, C8, by buffer exchanging AP180CTD-FL into 25 mM HEPES, 150 mM NaCl, 1 mM CdCl_2_, pH 7.35 through a 10^14^-fold dilution of the solution using Amicon 10K spin filter (MilliporeSigma cat#UFC501024) washes at 4C. Specifically, we diluted 50 μL of protein with 450 μL of buffer for a total 10x dilution and centrifuged the spin filter unit at 17,000 x g for 8 minutes to concentrate the solution back down to 50 μL. We repeated this dilution and concentration 6 more times for a total of 7 washes. We then transferred the washed protein solution to another spin filter unit and repeated the washes another 7 times for a total of 14 washes. After buffer exchange into the CdCl_2_ buffer, we incubated the protein with a 2-fold molar excess of ATTO 488-maleimide for 45 minutes at room temperature to label C159. The protein-dye mixture was then diluted 10-fold into 25 mM HEPES, 150 mM NaCl, 10mM EDTA pH 7.35 and incubated for 10 minutes at room temperature to deprotect C8 by chelating the CdCl_2_ in solution. We washed out the excess free dye, CdCl_2_, and EDTA with another 10^14^-fold dilution of the solution through 10 K Amicon spin filter washing using 25 mM HEPES, 150 mM NaCl, pH 7.35 at 4C. Finally, vesicles were prepared according to the “Lipid vesicle preparation for FRET” section, containing 73.5 mol % DOPC, 0.5 mol % DSPE-maleimide, 16 mol % Ni^2+^-DGS-NTA, and 10 mol % Texas Red-DHPE. The 488-labeled AP180CTD-FL was incubated with the vesicles for 1 hour at room temperature. The vesicle-protein mixture was then washed with 100 K Amicon spin filters and the excess, unconjugated protein was diluted out by 10^14^-fold. The resulting vesicles had N-terminally conjugated, C159 ATTO 488-labeled AP180CTD-FL to be used as a protein FRET sensor.

### Measuring Membrane Coverage by AP180CTD-FL and AP180CTD-1/3

We used a fluorescence intensity-based approach, which we have previously reported, to obtain a measurement of membrane coverage by AP180CTD-FL and AP180CTD-1/3 (33). Vesicles for these experiments were composed of 81.95 mol % DOPC, 1 mol % DP-EG10-biotin, 1 mol % PEG 10K-DSPE, 16 mol % of Ni^2+^-DGS-NTA, and 0.05 mol % Texas Red-DHPE. Lipid vesicles were prepared as described above through extrusion. Imaging wells were made by placing silicone gaskets onto ultraclean coverslips. Wells were coated for 20 min with biotinylated PLL−PEG. The biotinylated PLL−PEG was prepared according to a previously published protocol (35). After incubation, the well was washed repeatedly with a buffer containing 25 mM HEPES, 150 mM NaCl, and 1 mM TCEP pH 7.35. Neutravidin was added to the well at a final concentration of 0.2 mg/mL, incubated for 10 min, and washed out with HEPES buffer. The biotinylated vesicles were diluted to 2 μM and incubated in the well for 15 minutes to bind to the neutravidin. The excess, unbound vesicles were washed out with HEPES buffer. Protein was then incubated in the well to bind to the vesicles at various concentrations. The tethered vesicles were then imaged on a Zeiss spinning disk confocal microscope. At least 10 images were taken for each protein concentration. All images were cropped to the center 171 × 171 pixels of the 512 × 512 pixel field of view to account for uneven illumination at the edges. Fluorescence amplitudes of diffraction-limited puncta were obtained using the cmeAnalysis particle detection software in Matlab (36). This program detected individual vesicles by fitting two-dimensional Gaussian profiles to each puncta in the lipid fluorescence channel. We then used the centroids of the fluorescent puncta in the lipid channel to define the search region for fluorescent puncta in the protein channel. The dimension of the search region was three times the standard deviation of the Gaussian fit to the point spread function of our microscope. Using these detected fluorescent intensities, we estimated vesicle diameter from lipid fluorescence values by computing a scaling factor, which centered the median of the vesicle brightness distribution, prior to adding protein, to the number-weighted average vesicle diameter obtained from dynamic light scattering. We estimated the number of bound proteins on each vesicle by comparing brightness values in the protein channel to the brightness of a single molecule of ATTO 488 labeled hisENTH, a globular protein used for the purpose of single molecule calibration, which was measured using the same microscope settings. Membrane coverage was then estimated by dividing the total area occupied by the bound proteins by the approximate surface area of the vesicle, equation 7.

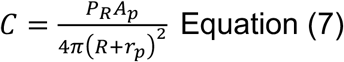

Here, C is total membrane coverage, P_R_ is number of proteins bound to the membrane, A_P_ is the area occupied on the membrane surface by the protein, R is the radius of the vesicle, and r_p_ is the radius of the protein. We approximated the area occupied per protein on the membrane, A_p_, by using hydrodynamic radius values, r_p_, previously reported by Zeno et al (16). Importantly, we also used equation 7 to calculate the membrane coverage by PEG 10K in the same manner, using an estimate of the radius of gyration (3.3 nm) as a freely-jointed random walk for PEG 10K. The total membrane coverage is the sum of the protein coverage and PEG coverage on the lipid vesicles.

### Crowding Lipid Vesicles With PEGylated lipids

To crowd the membrane surface with PEG chains of varying length, we used DSPE lipids with PEG groups covalently attached to the lipid head group by the manufacturer (Nanosoft Polymers). PEG 10K was included in all experiments as our polymeric sensor. For FRET experiments, PEG 10K-NH_2_ was labeled with ATTO 488 NHS ester. The crowding PEG lipids were left unlabeled. As described above, 10 mol% of Texas Red acceptor lipid was incorporated into the membrane. Separate batches of vesicles were created with an increasing mol % of crowding PEG lipid (2.25%, 3.5%, and 5%). For coverage experiments, the PEG 10K-NH_2_ sensor was left unlabeled, and instead we used a thiol group on the crowding PEGs to label with ATTO 488-maleimide. In this way, we keep the membrane composition the same while also being able to count the number of crowding PEGs without labeling complications from the PEG 10K sensor. Coverage experiments and analysis were performed as described above. To estimate the hydrodynamic radius of our various sizes of PEGs, we used a simple self-avoiding random walk model according to equation 8 (37):

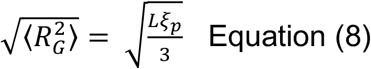

where 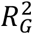 is radius of gyration, L is contour length of the polymer, and *ξ*_*p*_ is persistence length of the polymer. We used a previously measured experimental value of 3.8 Å for the persistence length of PEG to calculate 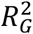 for our various PEG sizes (38).

### Measuring Membrane Fission by AP180CTD-FL and AP180CTD-1/3

We used the experimental methods described in the “Measuring Membrane Coverage by AP180CTD-FL and AP180CTD-1/3” section to measure vesicle size over a range of membrane coverages. Vesicles were composed of 81.95 mol % DOPC, 1 mol % DP-EG10-biotin, 1 mol % PEG 10K-DSPE, 16 mol % of Ni^2+^-DGS-NTA, and 0.05 mol % Texas Red-DHPE. Lipid vesicles were prepared through extrusion as described in the “Lipid vesicle preparation for FRET” section. AP180CTD-FL or AP180CTD-1/3 proteins at varying concentrations (20 nM to 4 μM) were incubated with vesicles for 1 hour at room temperature. The protein−vesicle solution was then tethered to the biotinylated PLL−PEG functionalized coverslip for 1 hour. Image acquisition and analysis was performed as described above. Notably, we estimated vesicle diameter from lipid fluorescence values by computing a scaling factor, which centered the median of the vesicle brightness distribution, prior to adding protein, to the number-weighted average vesicle diameter obtained from dynamic light scattering (DLS). This DLS measurement was performed for every set of vesicles on the day of the experiment. After addition of protein to the sample, the fluorescence intensity of each tethered vesicle puncta in the lipid channel decreased according to how much fission occurred and how small the vesicles became. This change in vesicle size over a range of membrane coverages by AP180CTD-FL or AP180CTD-1/3 shown in Figure 5E.

## Author Contributions

Justin R. Houser designed and performed experiments. Hyun Woo Cho provided key thermodynamic insight into the interpretation of data. Carl C. Hayden wrote custom software for data analysis and helped design experiments. Noel X. Yang helped with the preparation of materials and reagents and performed experiments. Liping Wang purified critical proteins for experiments. In addition, all authors consulted together on the interpretation of results and the preparation of the manuscript.

## Acknowledgements

This research was supported by the National Science Foundation through a Modulus Grant BIO-1934509 to J.C.S. and by CHE Grant 19-00093 to D.T. This work was also supported by the Welch Foundation through Grants F-2047 to J.C.S and F-0019 to D.T., by the Collie-Welch chair (D.T.), and by a grant from the National Institutes of Health (Grant R35GM139531) to J.C.S.

## References

1. Zeno, W. F., K. J. Day, V. D. Gordon, and J. C. Stachowiak. 2020. Principles and Applications of Biological Membrane Organization. Annu Rev Biophys. 49:19–39, doi: 10.1146/annurev-biophys-121219-081637.

2. Stachowiak, J. C., F. M. Brodsky, and E. A. Miller. 2013. A cost-benefit analysis of the physical mechanisms of membrane curvature. Nat Cell Biol. 15(9):1019–1027, doi: 10.1038/ncb2832.

3. Praefcke, G. J., and H. T. McMahon. 2004. The dynamin superfamily: universal membrane tubulation and fission molecules? Nat Rev Mol Cell Biol. 5(2):133–147, doi: 10.1038/nrm1313.

4. Jiang, Z., M. de Messieres, and J. C. Lee. 2013. Membrane remodeling by α-synuclein and effects on amyloid formation. J Am Chem Soc. 135(43):15970–15973, doi: 10.1021/ja405993r.

5. Shurer, C. R., J. C. Kuo, L. M. Roberts, J. G. Gandhi, M. J. Colville, T. A. Enoki, H. Pan, J. Su, J. M. Noble, M. J. Hollander, J. P. O’Donnell, R. Yin, K. Pedram, L. Möckl, L. F. Kourkoutis, W. E. Moerner, C. R. Bertozzi, G. W. Feigenson, H. L. Reesink, and M. J. Paszek. 2019. Physical Principles of Membrane Shape Regulation by the Glycocalyx. Cell. 177(7):1757-1770.e1721, doi: 10.1016/j.cell.2019.04.017.

6. Frost, A., V. M. Unger, and P. De Camilli. 2009. The BAR domain superfamily: membrane-molding macromolecules. Cell. 137(2):191–196, doi: 10.1016/j.cell.2009.04.010.

7. Fotin, A., Y. Cheng, P. Sliz, N. Grigorieff, S. C. Harrison, T. Kirchhausen, and T. Walz. 2004. Molecular model for a complete clathrin lattice from electron cryomicroscopy. Nature. 432(7017):573–579, doi: 10.1038/nature03079.

8. Kirchhausen, T. 2012. Bending membranes. Nat Cell Biol. 14(9):906–908, doi: 10.1038/ncb2570.

9. Busch, D. J., J. R. Houser, C. C. Hayden, M. B. Sherman, E. M. Lafer, and J. C. Stachowiak. 2015. Intrinsically disordered proteins drive membrane curvature. Nature Communications. 6(1):1–11, doi: 10.1038/ncomms8875.

10. Snead, W. T., W. F. Zeno, G. Kago, R. W. Perkins, J. B. Richter, C. Zhao, E. M. Lafer, and J. C. Stachowiak. 2019. BAR scaffolds drive membrane fission by crowding disordered domains. J Cell Biol. 218(2):664–682, doi: 10.1083/jcb.201807119.

11. Scheve, C. S., P. A. Gonzales, N. Momin, and J. C. Stachowiak. 2013. Steric pressure between membrane-bound proteins opposes lipid phase separation. J Am Chem Soc. 135(4):1185–1188, doi: 10.1021/ja3099867.

12. Singh, P., P. Mahata, T. Baumgart, and S. L. Das. 2012. Curvature sorting of proteins on a cylindrical lipid membrane tether connected to a reservoir. Phys Rev E Stat Nonlin Soft Matter Phys. 85(5 Pt 1):051906, doi: 10.1103/PhysRevE.85.051906.

13. Derganc, J., and A. Čopič. 2016. Membrane bending by protein crowding is affected by protein lateral confinement. Biochim Biophys Acta. 1858(6):1152–1159, doi: 10.1016/j.bbamem.2016.03.009.

14. Houser, J. R., C. C. Hayden, D. Thirumalai, and J. C. Stachowiak. 2020. A Förster Resonance Energy Transfer-Based Sensor of Steric Pressure on Membrane Surfaces. J Am Chem Soc. 142(49):20796–20805, doi: 10.1021/jacs.0c09802.

15. Morris, S. A., S. Schröder, U. Plessmann, K. Weber, and E. Ungewickell. 1993. Clathrin assembly protein AP180: primary structure, domain organization and identification of a clathrin binding site. EMBO J. 12(2):667–675. doi: 10.1038/emm.1999.31.

16. Zeno, W. F., A. S. Thatte, L. Wang, W. T. Snead, E. M. Lafer, and J. C. Stachowiak. 2019. Molecular Mechanisms of Membrane Curvature Sensing by a Disordered Protein. J Am Chem Soc. 141(26):10361–10371, doi: 10.1021/jacs.9b03927.

17. Brown, H. G., and J. H. Hoh. 1997. Entropic exclusion by neurofilament sidearms: a mechanism for maintaining interfilament spacing. Biochemistry. 36(49):15035–15040, doi: 10.1021/bi9721748.

18. Kumar, S., X. Yin, B. D. Trapp, J. H. Hoh, and M. E. Paulaitis. 2002. Relating interactions between neurofilaments to the structure of axonal neurofilament distributions through polymer brush models. Biophys J. 82(5):2360–2372, doi: 10.1016/S0006-3495(02)75581-1.

19. Szasz, C. S., A. Alexa, K. Toth, M. Rakacs, J. Langowski, and P. Tompa. 2011. Protein disorder prevails under crowded conditions. Biochemistry. 50(26):5834–5844, doi: 10.1021/bi200365j.

20. Hong, J., and L. Gierasch. 2010. Macromolecular Crowding Remodels the Energy Landscape of a Protein by Favoring a More Compact Unfolded State. J. Am. Chem. Soc. 132(30): 10445–10452. doi: 10.1021/ja103166y

21. Johansen, D., C. M. Jeffries, B. Hammouda, J. Trewhella, and D. P. Goldenberg. 2011. Effects of macromolecular crowding on an intrinsically disordered protein characterized by small-angle neutron scattering with contrast matching. Biophys J. 100(4):1120–1128, doi: 10.1016/j.bpj.2011.01.020.

22. Soranno, A., I. Koenig, M. B. Borgia, H. Hofmann, F. Zosel, D. Nettels, and B. Schuler. 2014. Single-molecule spectroscopy reveals polymer effects of disordered proteins in crowded environments. Proc Natl Acad Sci U S A. 111(13):4874–4879, doi: 10.1073/pnas.1322611111.

23. Kaufmann, S., O. Borisov, M. Textora, and E. Reimhult. 2011. Mechanical properties of mushroom and brush poly(ethylene glycol)-phospholipid membranes. Soft Matter. 7:9267–9275. doi: 10.1039/C1SM05746D

24. Lipowsky, R. 1995. Bending of membranes by anchored polymers. Europhys. Lett. 30(4):197–202. doi: 10.1209/0295-5075/30/4/002

25. Dimarzio, E. A., and F. L. McCrackin. 1965. One-dimensional model of polymer adsorption. The Journal of Chemical Physics. 43(2):539–547, doi: 10.1063/1.1696778.

26. Snead, W. T., and J. C. Stachowiak. 2018. A Tethered Vesicle Assay for High-Throughput Quantification of Membrane Fission. Methods Enzymol. 611:559–582, doi: 10.1016/bs.mie.2018.08.014.

27. Puljung, M. C., and W. N. Zagotta. 2011. Labeling of specific cysteines in proteins using reversible metal protection. Biophys J. 100(10):2513–2521, doi: 10.1016/j.bpj.2011.03.063.

28. Chen, W., R. Cordero, H. Tran, and C. K. Ober. 2017. 50th Anniversary Perspective: Polymer Brushes: Novel Surfaces for Future Materials. Macromolecules. 50(11):4089–4113. doi: 10.1021/acs.macromol.7b00450

29. de Gennes, P. G. 1980. Conformations of Polymers Attached to an Interface. Macromolecules. 13(5):1069–1075. doi:10.1021/ma60077a009

30. de Gennes, P. G. 1979. Scaling Concepts in Polymer Physics. Cornell University Press.

31. Carnahan, N. F., and K. E. Starling. 1969. Equation of state for nonattracting rigid spheres. The Journal of Chemical Physics. 51(2):635–636, doi: 10.1063/1.1672048.

32. des Cloizeaux, J. 1975. The Lagrangian theory of polymer solutions at intermediate concentrations. J. Phys. 36(4)281–291. doi:10.1051/jphys:01975003604028100.

33. Snead, W. T., C. C. Hayden, A. K. Gadok, C. Zhao, E. M. Lafer, P. Rangamani, and J. C. Stachowiak. 2017. Membrane fission by protein crowding. Proceedings of the National Academy of Sciences of the United States of America. 114(16):E3258–E3267, doi: 10.1073/pnas.1616199114.

34. Pietrosemoli, N., R. Pancsa, and P. Tompa. 2013. Structural disorder provides increased adaptability for vesicle trafficking pathways. PLoS Comput Biol. 9(7):e1003144, doi: 10.1371/journal.pcbi.1003144.

35. Kunding, A. H., M. W. Mortensen, S. M. Christensen, and D. Stamou. 2008. A fluorescence-based technique to construct size distributions from single-object measurements: application to the extrusion of lipid vesicles. Biophys J. 95(3):1176–1188, doi: 10.1529/biophysj.108.128819.

36. Aguet, F., C. N. Antonescu, M. Mettlen, S. L. Schmid, and G. Danuser. 2013. Advances in analysis of low signal-to-noise images link dynamin and AP2 to the functions of an endocytic checkpoint. Developmental Cell. 26(3):279–291, doi: 10.1016/j.devcel.2013.06.019.

37. Phillips, R., J. Kondev, J. Theriot, and H. G. Garcia. 2009. Physical Biology of the Cell. Garland Science.

38. Kienberger, F., V. P. Pastushenko, G. Kada, H. J. Gruber, C. Riener, H. Schindler, and P. Hinterdorfer. 2000. Static and Dynamical Properties of Single Poly(Ethylene Glycol) Molecules Investigated by Force Spectroscopy. Single Molecules. 1(2):123–128, doi: 10.1002/1438-5171(200006)1:2<123::AID-SIMO123>3.0.CO;2-3.

