## Supporting Information for "Molecular Mechanisms of Steric Pressure Generation and Membrane Remodeling by Intrinsically Disordered Proteins"

```
(Linker region)    |(AP180CTD start)
        10         20         30         40         50         60
GSPGIPHCPH ASSVPRVDIF ATASAAAPVS SAKPSSDLLD LQPDFSGAAA GAAAPVPPPT

        70         80         90        100        110        120
GGATAWGDLL GEDSLAALSS VPSEAPISDP FAPEPSPPTT TTEPASASAS ATTAVTAATT

       130        140        150        160        170        180
EVDLFGDAFA ASPGEAPAAS EGATAPATPA PVAAALDACS GNDPFAPSEG SAEAAPELDL

       190        200        210        220        230        240
FAMKPPETSA PVVTPTASTA PPVPATAPSP APTAVAATAA ATTTTTAAAA TTTATTSAAA

       250        260        270        280        290        300
AATAAAPPAL DIFGDLFDSA PEVAAASKPD VAPSIDLFGT DAFSSPPRGA SPVPESSLTA

       310        320        330        340        350        360
DLLSVDAFAA PSPASTASPA KAESSGVIDL FGDAFGSSAS ETQPAPQAVS SSSASADLLA

       370        380        390        400        410        420
GFGGSFMAPS TTPVTPAQNN LLQPNFEAAF GTTPSTSSSS SFDPSGDLLM PTMAPSGQPA

       430        440        450        460        470        480
PVSMVPPSPA MSASKGLGSD LDSSLASLVG NLGISGTTSK KGDLQWNAGE KKLTGGANWQ

       490        500        510        520        530        540
PKVTPATWSA GVPPQGTVPP TSSVPPGAGA PSVGQPGAGY GMPPAGTGMT MMPQQPVMFA

       550        560        570        580
QPMMRPPFGA AAVPGTQLSP SPTPATQSPK KPPAKDPLAD LNIKDFL
```

**Figure S1: Amino acid sequence for the AP180CTD sensor.** The amino acid sequence for AP180CTD used for the sensor, derived from AP180 in Rattus norvegicus (residues 328-898; CAA48748). The linker region contains a cysteine directly adjacent to a histidine used for conjugation techniques described in the methods section.

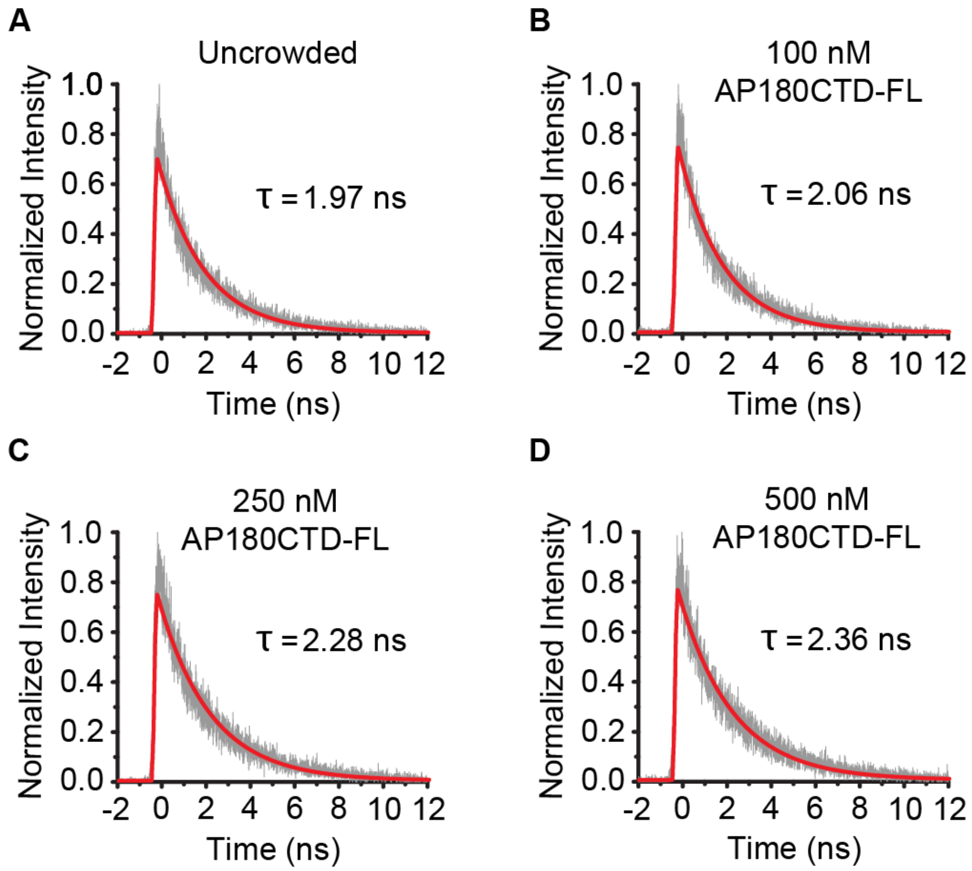


**Figure S2: Single exponential fits of AP180CTD sensor crowding data.** (A-D) Representative decay curves picked from data in Figure 2C, D and fit by a single component exponential curve. All curves were fit in Becker & Hickl SPCImage software using a weighted least squares method and convoluted with the instrument response function as determined from the signal from nonresonant Raman scattering in water. Mean lifetimes were calculated as the value of the 1/e single exponential fit.

AP180CTD    AP180CTD-1/3

```
(6x histag linker region) |(AP180CTD start)
        10         20         30         40         50         60
GSPGIPHHHH HHTDPHASSV PR|VDIFATAS AAAPVSSAKP SSDLLDLQPD FSGAAAGAAA

        70         80         90        100        110        120
PVPPPTGGAT AWGDLLGEDS LAALSSVPSE APISDPFAPE PSPPTTTTEP ASASASATTA

       130        140        150        160        170        180
VTAATTEVDL FGDAFAASPG EAPAASEGAT APATPAPVAA ALDACSGNDP FAPSEGSAEA
                         (1/3 end)|
       190        200        210        220        230        240
APELDLFAMK PPETSAPVVT PTASTAPPVP AT|APSPAPTA VAATAAATTT TTAAAATTTA

       250        260        270        280        290        300
TTSAAAAATA AAPPALDIFG DLFDSAPEVA AASKPDVAPS IDLFGTDAFS SPPRGASPVP

       310        320        330        340        350        360
ESSLTADLLS VDAFAAPSPA STASPAKAES SGVIDLFGDA FGSSASETQP APQAVSSSSA

       370        380        390        400        410        420
SADLLAGFGG SFMAPSTTPV TPAQNNLLQP NFEAAFGTTP STSSSSSFDP SGDLLMPTMA

       430        440        450        460        470        480
PSGQPAPVSM VPPSPAMSAS KGLGSDLDSS LASLVGNLGI SGTTSKKGDL QWNAGEKKLT

       490        500        510        520        530        540
GGANWQPKVT PATWSAGVPP QGTVPPTSSV PPGAGAPSVG QPGAGYGMPP AGTGMTMMPQ
                                             (AP180CTD end)|
       550        560        570        580        590
QPVMFAQPMM RPPFGAAAVP GTQLSPSPTP ATQSPKKPPA KDPLADLNIK DFL|
```

**Figure S3: Amino acid sequence for AP180CTD-FL and AP180CTD-1/3 truncation.** The amino acid sequences for AP180CTD-FL and its 1/3 truncation mutant derived from AP180 in Rattus norvegicus (residues 328-898; CAA48748). The first 22 amino acid residues correspond to the linker region, containing a 6x histidine tag.
